# OctopusV and TentacleSV: a one-stop toolkit for multi-sample, cross-platform structural variant comparison and analysis

**DOI:** 10.1101/2025.03.24.645012

**Authors:** Qingxiang Guo, Yangyang Li, Ting-You Wang, Abhi Ramakrishnan, Rendong Yang

## Abstract

Structural variants (SVs) significantly influence genomic variability and disease, but their accurate analysis across multiple samples and sequencing platforms remains challenging. We developed OctopusV, a tool that standardizes ambiguous breakend (BND) annotations into canonical SV types (inversions, duplications, translocations) and integrates variant calls using flexible set operations, such as union, intersection, difference, and complement, enabling cohort-specific variant identification. Together with TentacleSV, an automated pipeline, OctopusV provides an end-to-end solution from raw data to final callsets. Evaluations show improved precision, recall, and consistency, highlighting its value in cancer genomics and rare disease diagnostics. Both tools are available at https://github.com/ylab-hi/OctopusV and https://github.com/ylab-hi/TentacleSV.

## Background

Structural variants (SVs)—genomic alterations spanning tens to thousands of base pairs—greatly impact genomic architecture and function through changes in gene regulation, dosage modification, and chromosomal rearrangements [1–3]. These variants, which include deletions (DELs), insertions (INSs), inversions (INVs), duplications (DUPs), and translocations (TRAs), are essential in cancer genomics and rare disease research where their accurate detection directly affects clinical diagnosis and treatment [5,6]. The rapid growth of sequencing technologies, such as short-read (Illumina) and long-read (PacBio, Oxford Nanopore) platforms, has produced extensive genomic datasets, creating an urgent need for computational frameworks that can reliably analyze SVs across diverse biological contexts and technological platforms [6–8].

The scientific community has developed numerous tools to address SV detection and integration [9]. For instance, SURVIVOR [6] enables SV merging across multiple samples using positional overlap, Jasmine [10] refines breakpoint locations through clustering techniques, SVmerge [11] and CombiSV [12] support population-level analyses by combining calls from multiple samples. Viola-SV [13] provides Python-based manipulation and visualization of SV callsets, whereas general-purpose tools such as BCFtools [14] and VCFtools [15] primarily handle SNP/indel operations without specialized functions for complex SV merging. Additionally, PanPop [16] effectively merges SVs at the population scale with a sequence-aware approach, but primarily addresses large-scale genotyping and allele frequency estimation rather than specialized SV merging strategies.

Despite these advances, several critical challenges remain in SV analysis, particularly for applications in precision medicine and large-scale genomic studies [17]. First, widely-used SV detection tools—including Manta [18], LUMPY [19], and SvABA [20]—frequently report breakend (BND) annotations [2], which describe breakpoints without specifying the precise SV type. Without proper handling, BND events are often discarded [21] or misclassified, potentially omitting biologically and clinically relevant variants [22]. Moreover, different SV callers may represent identical variants through different conventions (e.g., reporting an inversion as either INV or as ambiguous BND events), making accurate merging difficult and reducing overall analysis reliability [23]. Second, existing SV merging tools mainly support basic operations such as union or intersection, which are not sufficient for the detailed analyses needed in modern genomic research [24]. For example, cancer investigations often require identification of tumor-specific SVs or variants unique to specific cancer subtypes, operations that currently require custom scripting and lack standardized solutions. This limitation creates barriers for researchers without extensive programming expertise. Third, the SV analysis pipeline remains highly fragmented, requiring researchers to manage multiple independent software components, configure various parameters, and handle intermediate file formats [24]. This workflow complexity creates reproducibility challenges and limits scalability, especially for large-scale sequencing projects and clinical applications [3,25].

To address these fundamental challenges, we have developed OctopusV, a versatile toolkit that enhances SV analysis by integrating variant calls from multiple detection methods and sequencing platforms. Together with TentacleSV, an automated pipeline, our toolkit provides an end-to-end workflow from raw sequencing data to high-confidence SV calls. OctopusV advances beyond existing tools through three key innovations. First, it features a specialized BND correction module that systematically converts ambiguous BND annotations into canonical SV types (INV, DUP, TRA). This not only recovers biologically important variants that would otherwise be discarded but also enhances merging accuracy. Our evaluations show that this approach delivers measurable improvements in precision, recall, SV type consistency, and overall merging accuracy across diverse datasets compared to current methods. Second, OctopusV introduces advanced set operations, including difference, complement, and user-defined custom operations that enable sophisticated variant filtering strategies (e.g., identifying SVs unique to specific sample groups), eliminating the need for manual scripting. Third, TentacleSV automates the entire SV analysis process with minimal intervention, ensuring consistent analyses across projects. Additional features include benchmarking against reference datasets and visualization options (e.g., SV type distributions, chromosome maps), which facilitate result interpretation. Currently, no other framework combines automated BND correction, flexible custom set merging, and a fully automated pipeline, making our approach potentially valuable for various applications such as cancer subtype-specific SV identification or cohort analyses [26].

## Results

### Overview of OctopusV architecture

OctopusV addresses two key challenges in SV analysis: standardizing ambiguous breakend (BND) annotations and integrating multi-caller outputs through flexible operations. As shown in **Fig. 1**, the software features three primary modules—the input processing layer, the BND correction module, and the SV merging module—along with additional functionalities for benchmarking, visualization, and format conversion. The input layer handles variant call format (VCF) files from multiple sequencing platforms, including next-generation sequencing (NGS), Pacific Biosciences (PacBio), and Oxford Nanopore Technologies (ONT), and commonly used SV callers, such as Manta, LUMPY, SvABA, DELLY [27], PBSV [28], SVIM [29], Sniffles [6], CuteSV [30], SVDSS [31], and DeBreak [32], converting them into a standardized format for analysis. The BND correction module transforms BND events into canonical SV types, using pattern recognition and coordinate analysis, while the SV merging module integrates multi-caller outputs with strategies including intersection, union, support-based thresholds, custom set operations, and single-caller extractions. Additional features, such as the ‘stat’ and ‘plot’ subcommands, generate detailed SV metrics and publication-ready visualizations (e.g., UpSet plots, SV size distributions, chromosome maps), and a HTML reporting system enables result exploration (**Supplementary Fig. 1**).

**Fig. 1.**
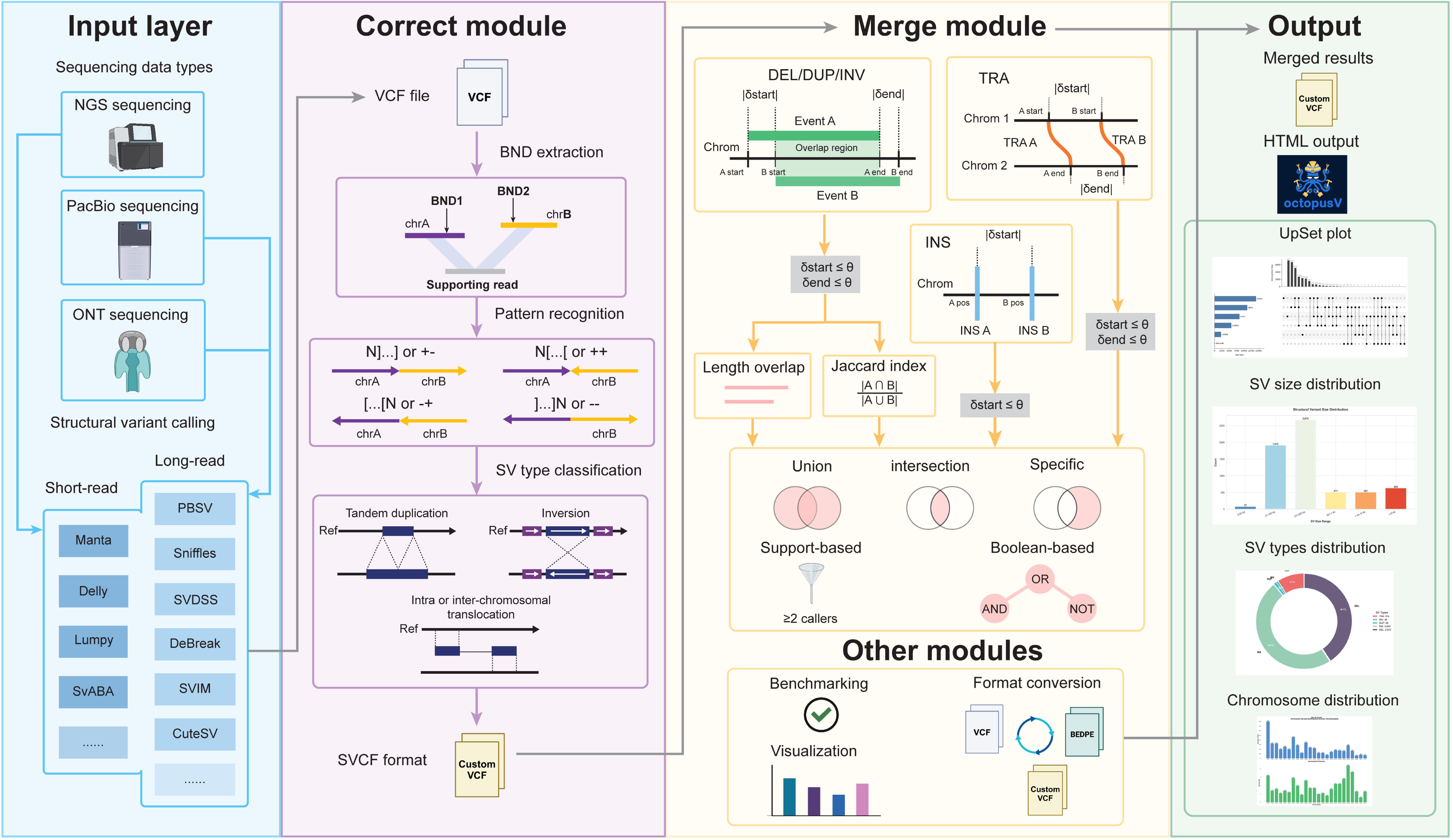
Schematic overview of OctopusV’s architecture for structural variants (SVs) standardization and merging. OctopusV is organized into three primary modules: the Input layer, the Correct module, and the Merge module, along with supporting functionality modules. The Input layer processes SV data from multiple sequencing platforms (NGS, PacBio, ONT, and others) and handles VCF files from standard SV callers. Several common callers are presented as examples; however, OctopusV can work with any VCF-formatted SV calls regardless of the caller or sequencing technology. The Correct module processes VCF files through sequential steps: BND extraction, pattern recognition of breakpoint orientations (shown by N[…] and […]N patterns), and SV type classification into standard forms (duplication, inversion, intra/inter-chromosomal TRA). The Merge module implements advanced merging strategies based on event coordinates and properties, supporting various operations including length overlap assessment, Jaccard index calculation, and set operations (union, intersection, specific). This module handles different SV types (DEL/DUP/INV, TRA, INS) with specific coordinate matching criteria (δstart, δend thresholds). Additional modules provide benchmarking, format conversion, and visualization capabilities. The output includes merged results in a customized VCF format and comprehensive visualization options (UpSet plots, SV size distribution, SV type distribution, and chromosome distribution). The customized VCF format serves as an intermediate representation that facilitates integration between modules. Additionally, OctopusV generates interactive HTML outputs that allow users to visualize and explore SV data through a web browser interface.

To enhance OctopusV’s adoption in research workflows, TentacleSV was developed as an automated Snakemake-based pipeline, streamlining the entire SV analysis process from raw sequencing data to final merged variants (**Supplementary Fig. 2**). The pipeline coordinates read mapping, multi-caller variant detection, and OctopusV’s BND correction and merging operations through a simple configuration file (config.yaml), requiring only user-specified inputs and parameters. TentacleSV simplifies workflow management, reduces manual intervention, and ensures compatibility with OctopusV’s core functionalities, enhancing reproducibility across large-scale analyses [21,24].

### Feature comparison with existing tools

To contextualize OctopusV’s capabilities among existing tools, we conducted feature comparison against four widely-used SV merging solutions: Jasmine, SURVIVOR, SVmerge, and CombiSV (**Fig. 2**). While several tools, such as Jasmine and SURVIVOR, offer partial support for flexible merging or format conversion, OctopusV provides an integrated solution that includes native BND correction, benchmarking, and custom set operations. Notably, **Fig. 2b** showcases OctopusV’s advanced set operation capabilities, enabling custom set operations (e.g., (Sample1, 2) only, (Sample1, 2, 3) only) for precise variant filtering across multiple samples—functionality not available in competing tools, which are typically restricted to basic union or intersection operations.

**Fig. 2.**
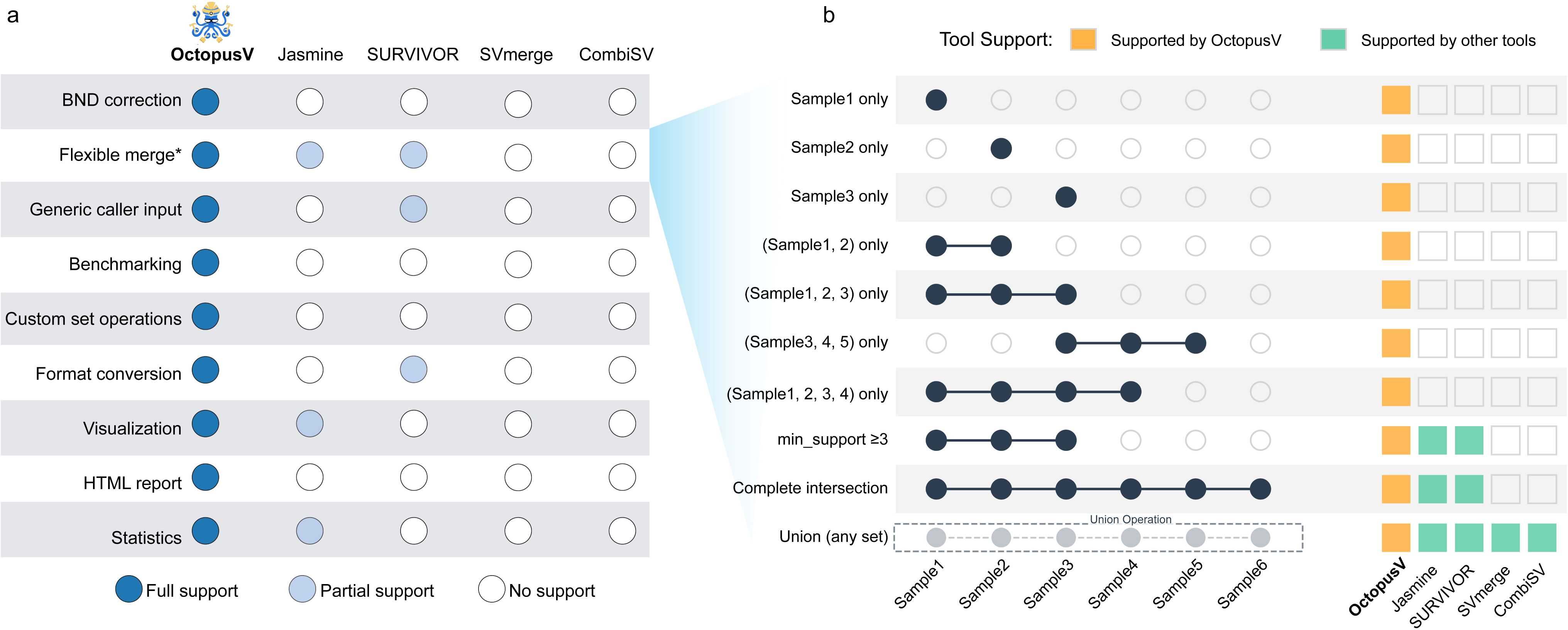
Comparison of key features between OctopusV and existing structural variants (SVs) merging tools (Jasmine, SURVIVOR, SVmerge, and CombiSV). **a** Feature comparison matrix showing support levels (full, partial, or none) for key functionalities across tools. **b** Advanced set operations supported by different tools, highlighting OctopusV’s flexible merge capabilities that enable custom set operations on SV sets from multiple samples. This functionality allows for the identification of sample-specific SVs through customizable set operations.

### BND prevalence and correction performance

To establish the importance of BND standardization in SV analysis, we first quantified the prevalence of BND annotations across different SV callers and sequencing platforms. Analyzing NA12878 [33] and VISOR [34] simulated datasets revealed considerable variation in BND usage among different tools (**Fig. 3a**). In the NA12878 NGS dataset, BND annotations constituted a considerable portion of SV calls, with SvABA reporting 100% of variants as BNDs, while Manta and LUMPY showed 62.11% and 67.93% BND representation, respectively. DELLY exhibited a more moderate BND usage at 17.40%. Analysis of the NA12878 long-read dataset revealed that long-read SV callers utilized BND annotations less frequently, though still at notable levels, ranging from 1.40% (PBSV) to 9.22% (CuteSV) in PacBio data. VISOR simulated datasets exhibited similar patterns, with SvABA again reporting 100% BND usage, Manta at 43.87%, and LUMPY at 11.00%. These findings confirm the widespread application of BND annotations across both real and simulated datasets.

**Fig. 3.**
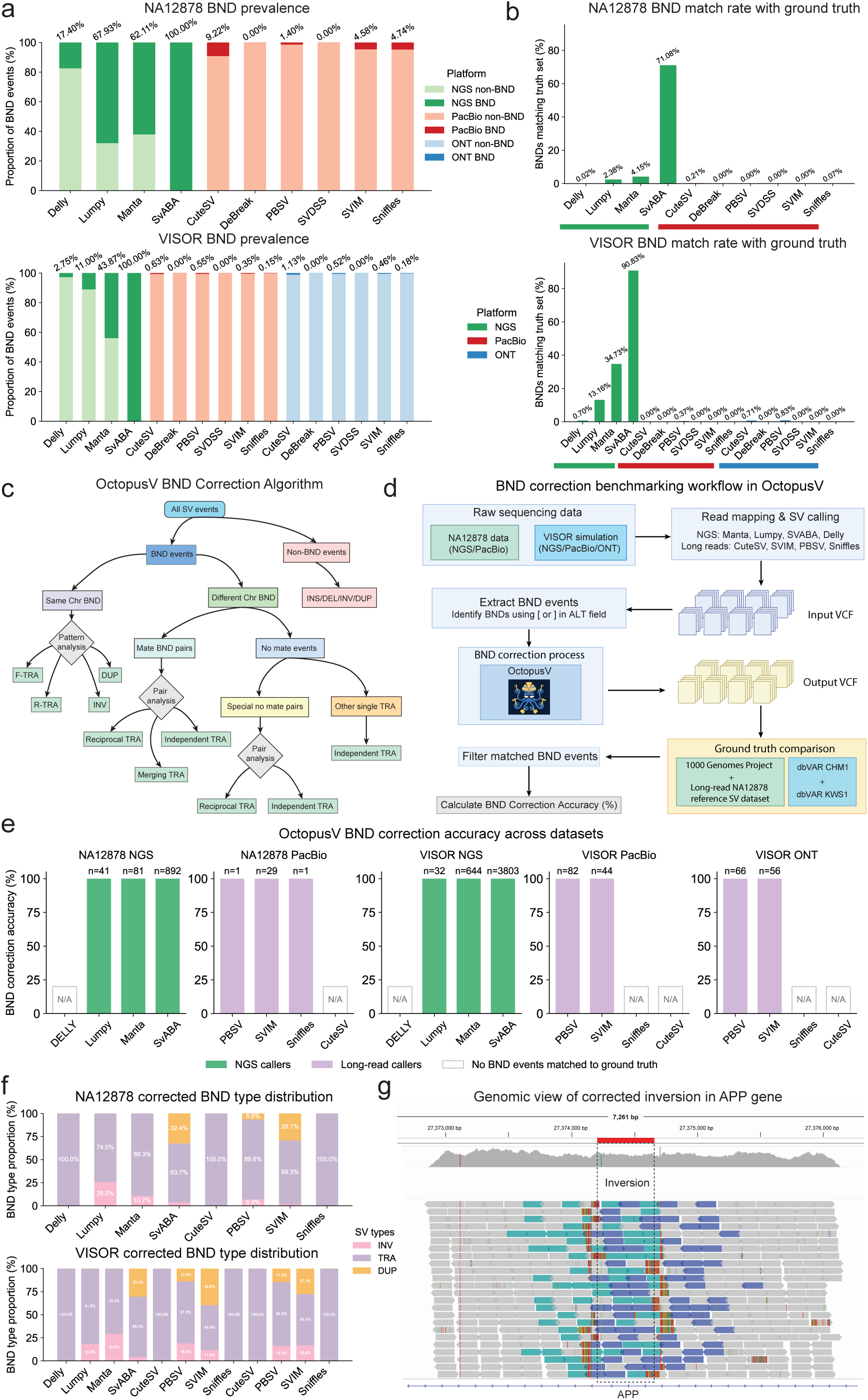
BND prevalence, correction, and benchmarking in OctopusV. **a** Proportion of BND annotations among total structural variants (SVs) reported by different callers in real (NA12878) and simulated (VISOR) datasets, highlighting frequent usage of BND annotations, particularly in short-read callers. **b** Proportion of BND calls matching high-confidence reference SV datasets, showing that a significant subset corresponds to genuine SVs. **c** OctopusV’s workflow for correcting ambiguous BND annotations by classifying them into canonical SV types (INV, DUP, TRA), based on chromosomal context and breakpoint patterns. **d** Benchmarking workflow for evaluating OctopusV’s correction accuracy through comparison with established ground-truth datasets. **e** Correction accuracy of OctopusV for short-read (purple bars) and long-read (blue bars) callers across datasets; white boxes indicate callers without matched BND events for correction evaluation. **f** Distribution of corrected BND events by canonical SV types (INV, DUP, TRA), revealing prevalent reclassification patterns across callers and sequencing platforms. **g** Example genomic view of a corrected inversion event in the APP gene from NA12878, initially identified as ambiguous BND events by Manta but resolved by OctopusV.

After quantifying BND prevalence, we next assessed their validity by comparing them against high-confidence reference sets (**Fig. 3b**). The match rate varied considerably by caller and platform. In NA12878 data, SvABA showed the highest correspondence at 71.08%, while other short-read callers demonstrated lower matching rates (4.15% for Manta, 2.38% for LUMPY, and 0.02% for DELLY). In VISOR simulated data, SvABA again exhibited the highest match rate (90.83%), followed by Manta (34.73%) and LUMPY (13.16%). The variation in match rates likely reflects algorithmic differences between callers and varying reference set comprehensiveness. However, these results indicate many BND events correspond to genuine SVs that would remain undetected without proper processing.

The BND correction module in OctopusV implements a hierarchical classification system to standardize SV representations (**Fig. 3c**). This system first categorizes events based on chromosomal context (intra-chromosomal versus inter-chromosomal), then analyzes breakpoint orientation patterns to determine specific variant types. To evaluate the correction accuracy, we established a benchmarking framework that independently processed BND events from each SV caller in both simulated datasets (VISOR NGS/ONT/PacBio) and real NA12878 data, with results validated against corresponding ground truth datasets (**Fig. 3d**). We focused solely on BND events that matched entries in the ground truth datasets. OctopusV correctly converted all these matched BND notations to their appropriate SV types across all tested configurations (**Fig. 3e**). We excluded from our assessment the BND calls from several callers (Sniffles and CuteSV in VISOR ONT and PacBio datasets, DELLY in VISOR NGS, and CuteSV in NA12878 PacBio) that did not match any ground truth variants. This approach allowed us to specifically evaluate OctopusV’s ability to standardize BND annotations without being confounded by SV caller performance variability. The consistent conversion accuracy observed across diverse platforms and callers indicates that OctopusV effectively standardizes BND annotations for validated SVs.

Analysis of the distribution of corrected BND events across final SV types (**Fig. 3f**) revealed distinct patterns specific to each caller and sequencing platform. In NA12878 data, DELLY-reported BNDs were exclusively reclassified as TRAs (100%), while LUMPY’s BNDs were predominantly converted to TRAs (73.97%) with the remainder as INVs (26.03%). Manta showed a similar pattern with 89.27% TRAs and 10.73% INVs. SvABA displayed the most diverse distribution, with 63.74% TRAs, 32.40% DUPs, and 3.86% INVs. Long-read callers followed comparable patterns, with most showing a predominance of TRA events. These distribution patterns were largely consistent between real and simulated datasets, with minor variations reflecting differences in underlying variant profiles.

To demonstrate the biological relevance of BND correction, we identified genes containing SVs from Manta-called BND events in NA12878 that were successfully corrected by OctopusV and subsequently validated against reference datasets. For example, an inversion affecting APP, a gene implicated in Alzheimer’s disease [35], was initially reported by Manta as ambiguous BND events. However, OctopusV correctly classified it as INV, which is further confirmed by AnnotSV annotation and manual inspection using IGV (**Fig. 3g**). Variants in other disease-associated genes were similarly identified, including a 12.5 kb inversion in the autism-associated CNTNAP2 gene [36] and a 4.8 kb inversion in SMC1A (linked to rare disease Cornelia de Lange syndrome) [5] (**Supplementary Dataset 1**). These results demonstrate OctopusV’s effectiveness in standardizing BND annotations, thereby enabling accurate identification of biologically relevant SVs.

### Evaluation of OctopusV merging performance across diverse datasets

To evaluate the merging capabilities of OctopusV, we compared its performance with four established tools—Jasmine, SURVIVOR, SVmerge, and CombiSV—using real (NA12878 NGS and PacBio) and simulated (VISOR NGS, PacBio, and ONT) datasets (**Fig. 4a; Supplementary Dataset 2)**. Performance metrics, including precision, recall, and F1-score, were assessed under common strategies (intersection, union, minimum support thresholds) and OctopusV-specific operations (maximum support, caller-specific extraction, and custom set operations) (**Fig. 4b**).

**Fig. 4.**
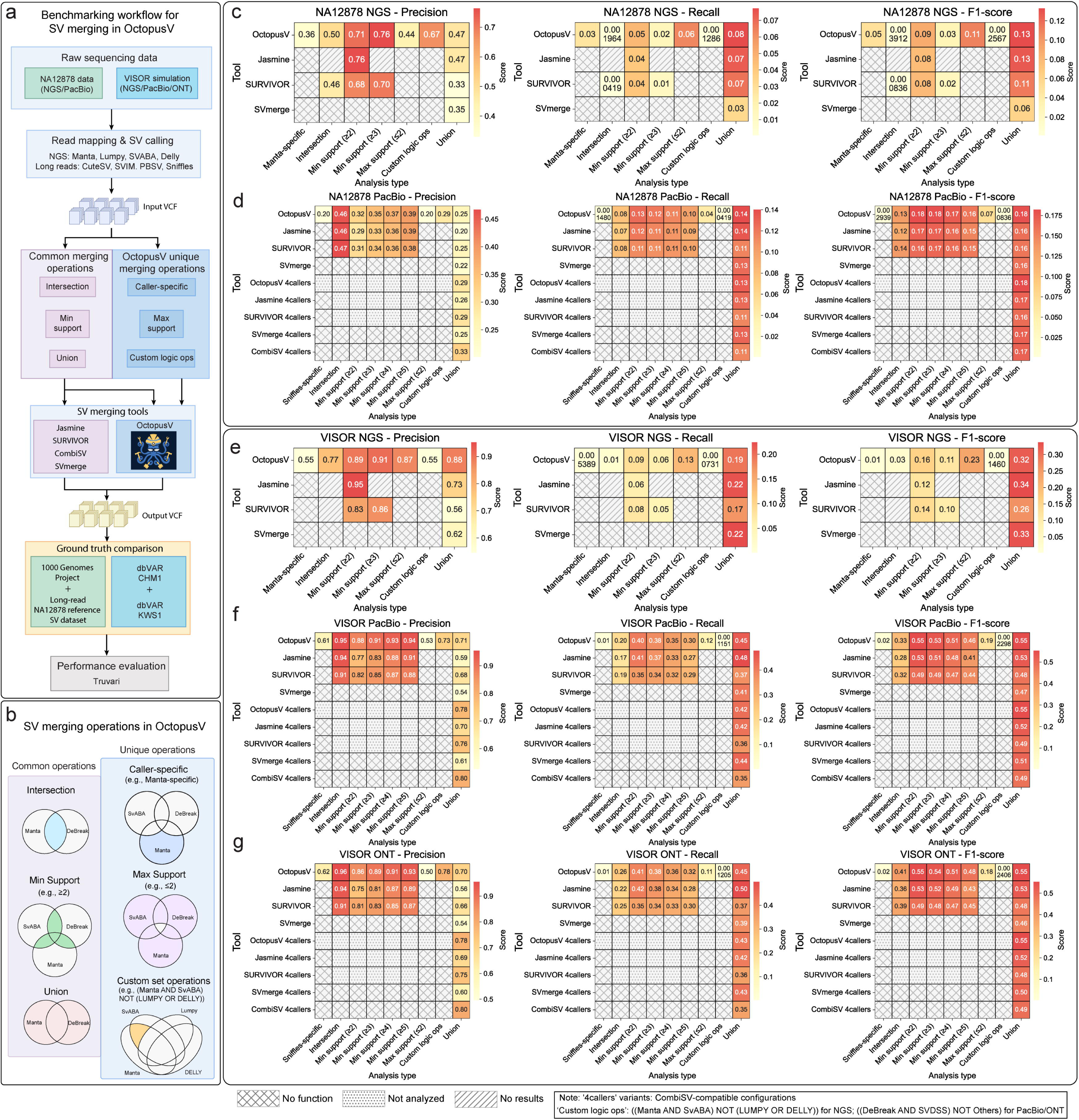
Evaluation of OctopusV’s SV merging strategies and performance compared to existing tools. **a** Workflow for benchmarking SV merging, illustrating processes from raw sequencing data to performance evaluation against reference datasets. The workflow includes SV calling to generate input VCF files, applying different merging strategies, and evaluating merged variants using Truvari. **b** Diagram illustrating SV merging operations supported by OctopusV. Common operations include intersection, union, and minimum support (variants supported by ≥2 callers). Unique operations provided by OctopusV include maximum support (variants supported by ≤2 callers), caller-specific extraction (e.g., Manta-only), and custom set logic operations. For NGS datasets (NA12878 NGS, VISOR NGS), the custom set logic operation is defined as ((Manta AND SvABA) NOT (LUMPY OR DELLY)). For PacBio and ONT datasets (NA12878 PacBio, VISOR PacBio, VISOR ONT), it is defined as ((DeBreak AND SVDSS) NOT Others). **c–g** Heatmaps comparing absolute Precision, Recall, and F1 scores of OctopusV against other SV merging tools (Jasmine, SURVIVOR, SVmerge, CombiSV) across multiple datasets: **c** NA12878 NGS, **d** NA12878 PacBio, **e** VISOR NGS, **f** VISOR PacBio, and **g** VISOR ONT. Cells marked with horizontal lines indicate unsupported operations; diagonal lines represent analyses with no results; dotted patterns indicate configurations not analyzed. The label "4callers" indicates analyses limited to the caller combination (Sniffles, PBSV, CuteSV, SVIM) specifically supported by CombiSV.

In the NA12878 NGS dataset (**Fig. 4c**), OctopusV showed competitive performance across standard merging strategies. The intersection strategy yielded precision of 0.50 and recall of 0.002, slightly surpassing SURVIVOR (precision 0.46, recall <0.001), while Jasmine did not produce results for this strategy. With minimum support thresholds, OctopusV demonstrated strong precision: 0.71 (≥2 callers) and 0.76 (≥3 callers). This performance was comparable to Jasmine’s precision of 0.76 for ≥2 callers (Jasmine provided no results for ≥3 callers) and exceeded SURVIVOR’s precision values of 0.68 (≥2 callers) and 0.70 (≥3 callers). However, recall values remained consistently low across all tools (OctopusV: 0.047 for ≥2 callers; Jasmine: 0.041 for ≥2 callers; SURVIVOR: 0.042 for ≥2 callers). Under the union strategy, OctopusV improved recall to 0.077, slightly above Jasmine (0.073), and more notably higher than SURVIVOR (0.068) and SVmerge (0.033), suggesting better integration of diverse caller outputs. For the NA12878 PacBio dataset (**Fig. 4d**), OctopusV demonstrated stable performance across merging strategies. The intersection strategy showed precision of 0.46 and recall of 0.077, closely aligning with SURVIVOR (0.47 and 0.080). Minimum support thresholds highlighted OctopusV’s strength, with precision increasing from 0.32 (≥2 callers) to 0.39 (≥5 callers) and recall stabilizing around 0.10, performing comparably to Jasmine (precision up to 0.39) and SURVIVOR (precision up to 0.38) across thresholds. The union strategy achieved an F1-score of 0.18 (precision 0.25, recall 0.14), exceeding Jasmine (0.16), SURVIVOR (0.16), and SVmerge (0.16). In a 4-caller subset (Sniffles, PBSV, CuteSV, SVIM), OctopusV’s F1-score reached 0.18 (precision 0.29, recall 0.13), surpassing CombiSV (0.17) and other tools.

In simulated datasets, OctopusV exhibited reliable performance. For VISOR NGS (**Fig. 4e**), the intersection strategy yielded precision of 0.77 and recall of 0.01, uniquely identifying intersecting variants that both Jasmine and SURVIVOR showed no results. This highlights OctopusV’s capability to capture critical variant intersections not reported by other tools. Minimum support thresholds showed OctopusV’s precision rising from 0.89 (≥2 callers) to 0.91 (≥3 callers) with recall peaking at 0.06, outperforming Jasmine (precision up to 0.95, recall 0.06) and SVmerge (precision up to 0.86, recall 0.05). The union strategy delivered an F1-score of 0.32 (precision 0.88, recall 0.19), competitive with Jasmine (0.34) and SVmerge (0.33). For VISOR PacBio (**Fig. 4f**), intersection precision was 0.95 with recall of 0.20, while minimum support peaked at 0.94 (≥5 callers) with recall at 0.30, surpassing Jasmine (precision up to 0.91, recall 0.27) and SURVIVOR (precision up to 0.88, recall 0.29). The union strategy achieved an F1-score of 0.55 (precision 0.71, recall 0.45). In VISOR ONT (**Fig. 4g**), intersection precision was 0.96 with recall of 0.26, minimum support reached 0.93 (≥5 callers) with recall of 0.32, and the union strategy yielded an F1-score of 0.55 (precision 0.70, recall 0.45), outperforming Jasmine (F1-score 0.53) and SVmerge (0.46). The 4-caller analyses across VISOR datasets maintained competitive F1-scores, with OctopusV reaching 0.55 (VISOR PacBio) and 0.55 (VISOR ONT), slightly exceeding CombiSV (0.49 and 0.49).

Low recall values were consistent across all tools, as seen in examples like 0.077 in NA12878 PacBio and 0.01 in VISOR NGS. This is likely due to variability in caller outputs, strict matching criteria, and incomplete ground truth datasets, particularly in complex genomic regions. Despite this, OctopusV’s advanced operations offered flexibility. Maximum support (≤2 callers) reached precision of 0.87 in VISOR NGS, caller-specific extraction (e.g., Manta for NGS, Sniffles for long-read) hit 0.62 in VISOR ONT, and custom set operations, e.g., ((DeBreak AND SVDSS) NOT Others), achieved 0.78 in VISOR ONT. These specialized merges help researchers focus on selected variants—such as highly confident ones or those found by specific callers—enabling more targeted analyses.

### SV type consistency and preservation

We evaluated SV type consistency and preservation across five merging tools—OctopusV, SURVIVOR, Jasmine, SVmerge, and CombiSV—using the same datasets as in the merging performance analysis (NA12878 NGS, NA12878 PacBio, VISOR NGS, VISOR ONT, and VISOR PacBio; **Fig. 5** and **Supplementary Fig. 3**). This analysis revealed distinct differences in variant type fidelity during merging.

**Fig. 5.**
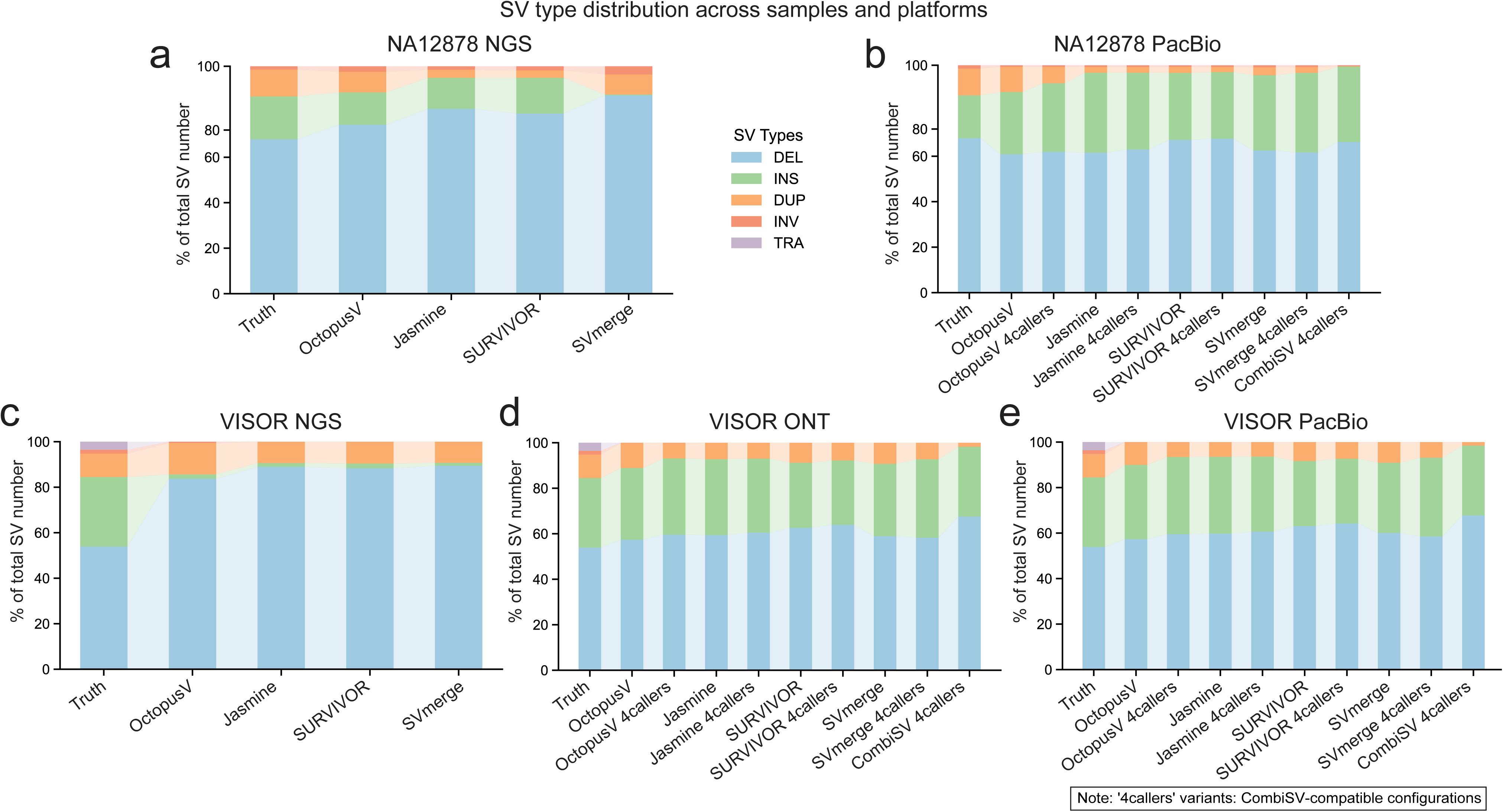
Preservation of SV type distribution for true positive SVs across merging tools and datasets. Normalized stacked bar plots comparing the distributions of true positive structural variants (SVs) identified by different merging tools (OctopusV, SURVIVOR, Jasmine, SVmerge, CombiSV) across five datasets: **a** NA12878 NGS, **b** NA12878 PacBio, **c** VISOR NGS, **d** VISOR ONT, and **e** VISOR PacBio. Each subplot specifically compares the SV type distribution of correctly identified variants (true positives) obtained by each tool with the ground truth distribution, illustrating each merging tool’s fidelity and consistency in preserving the original SV type annotations.

SURVIVOR showed substantial inconsistencies, with incorrect merges ranging from 183 instances in VISOR PacBio to 2,477 in VISOR NGS data (**Supplementary Fig. 3a**). These errors involved mismatches where the primary SVTYPE annotation differed from the original types of constituent variants, such as merging INV into DEL or DUP into TRA. Sankey diagrams illustrated these SV type annotation changes (**Supplementary Figs. 3b–f**), showing frequently observed SV type reclassifications (e.g., INV to DEL, DEL to INV) in SURVIVOR’s outputs. This included 935 INV-to-DEL misclassifications in NA12878 NGS (**Supplementary Fig. 3b**) and 2,004 DEL-to-INV conversions in VISOR NGS (**Supplementary Fig. 3d**). In contrast, OctopusV, Jasmine, SVmerge, and CombiSV reported zero incorrect merges across all datasets.

We also assessed how each tool preserved the original distribution of SV types compared to ground truth, focusing on true positive calls in union mode (**Figs. 5a–e**). In NA12878 NGS (**Fig. 5**), OctopusV’s type distribution closely aligned with ground truth, with DEL at 74.2%, INS at 20.6%, and DUP at 12.8%, showing minimal deviation, while other tools exhibited greater variability, such as SURVIVOR (DEL 72.3%, INS 20.3%, DUP 4.2%) and Jasmine (DEL 73.8%, INS 20.1%, DUP 5.2%). In NA12878 PacBio (**Fig. 5b**), OctopusV maintained DEL at 64.3%, INS at 23.1%, and DUP at 9.3%, closely matching ground truth (DEL 63.8%, INS 22.3%, DUP 11.1%), compared to SURVIVOR (DEL 65.1%, INS 20.4%, DUP 3.9%) and CombiSV (DEL 64.7%, INS 21.3%, DUP 0.1%).

In simulated datasets, OctopusV showed consistent fidelity. For VISOR NGS (**Fig. 5c**), OctopusV preserved DEL at 62.5%, DUP at 20.6%, and INS at 14.3%, closely tracking ground truth (DEL 62.1%, DUP 11.9%, INS 35.2%), while other tools deviated, such as SURVIVOR (DEL 60.3%, DUP 6.5%, INS 32.1%) and Jasmine (DEL 61.8%, DUP 10.1%, INS 27.9%).

Similar trends were observed in VISOR ONT (**Fig. 5d**) and VISOR PacBio (**Fig. 5e**), where OctopusV maintained DEL at approximately 60.6%, DUP at 17.8%, and INS at 20.7%, aligning closely with ground truth, while SURVIVOR and other tools showed greater deviations in DUP and INS proportions.

## Discussion

Accurate detection and integration of SVs across diverse samples, callers, and sequencing platforms remain a significant challenge for genomic research and clinical diagnostics. In this study, we introduced OctopusV and TentacleSV to address three key issues: (i) standardizing ambiguous BND annotations, (ii) enabling flexible SV merging strategies, and (iii) automating the SV analysis workflow. By converting BNDs into canonical SV types (INV, DUP, TRA), OctopusV helps recover potentially important variants that might otherwise be discarded as uncharacterized breakpoints [37]. Our benchmarks demonstrated that these standardized conversions enhance the interpretability of multi-caller SV callsets and provide a more consistent foundation for downstream analyses.

A central feature of OctopusV is its ability to perform advanced set operations, including difference and complement merges. While basic merging strategies (e.g., intersection, union, minimum support) are adequate for many routine analyses, OctopusV extends these capabilities by allowing difference and complement operations. These additional functions provide researchers with finer control over SV callsets, particularly valuable in various research contexts. For example, in clinical applications, these operations facilitate the identification of tumor-specific SVs compared to matched controls. At the technical level, custom set operations such as ((Manta AND SvABA) NOT (LUMPY OR DELLY)) and ((DeBreak AND SVDSS) NOT Others) enable targeted investigation of SV caller characteristics and algorithmic preferences, which can guide caller selection and pipeline optimization. Similarly, OctopusV’s maximum support functionality (variants supported by ≤2 callers/samples) helps researchers investigate low-frequency or low-confidence SVs, which may represent either artifacts or biologically relevant rare variants that are typically filtered out by conventional merging approaches. In addition, OctopusV preserves SV type fidelity during merging, reducing errors introduced by misclassification of variant types—a common issue in existing tools. Given that different SV types have distinct biological consequences, maintaining accurate annotations is essential for downstream functional analysis and clinical interpretation. By integrating these capabilities into TentacleSV, our Snakemake-based pipeline, we streamline the entire SV analysis workflow, ensuring consistency and reproducibility across large-scale or clinical genomic studies [38].

Despite these advantages, several limitations remain. First, the accuracy of OctopusV’s merging results depends on the quality of upstream SV callers, and biases or errors in the original variant calls cannot always be corrected through merging alone [39]. Second, SVs located in repetitive or highly complex genomic regions (e.g., centromeres, telomeric repeats, and assembly gaps) remain difficult to resolve, often yielding inconsistent calls across different SV callers [12,40]. Incorporating region-specific filters based on repeat annotations or developing confidence scoring approaches could help mitigate these challenges. Third, the limited availability of high-confidence SV datasets, especially for complex rearrangements, remains a challenge for benchmarking SV analysis tools [11], including OctopusV. Expanding reference datasets and improving ground truth annotations will be crucial for further refining performance assessments. Finally, while we have demonstrated the feasibility of custom set operations in various scenarios, further clinical validation will be crucial for translating these strategies into precision medicine applications.

OctopusV and TentacleSV serve as a bridge between experimental biologists and computational analysts. This connection enables researchers without specialized programming skills to perform sophisticated SV analyses. Our tools facilitate diverse applications including tumor-specific variant detection in cancer studies, identification of pathogenic SVs in rare disease diagnosis, and comparison of structural variant patterns across populations. Through flexible set operations that go beyond simple unions and intersections, researchers can define custom variant relationships and extract biologically meaningful patterns from complex genomic datasets. This improved analytical capability helps biologists better understand how structural variants contribute to phenotypic variation and disease mechanisms.

Future efforts will focus on enhancing variant prioritization, incorporating functional annotations, and extending custom set operations to enable annotation-based filtering. Additionally, machine learning [41] approaches could be explored for dynamically optimizing merging parameters. Integration with pan-genome frameworks will support population-scale analyses, while enhanced visualization capabilities will further simplify result interpretation for non-specialists. These developments will further support SV biomarker discovery and improve the utility of SV analysis in precision medicine and genomic diagnostics.

## Conclusions

OctopusV and TentacleSV provide an integrated framework addressing persistent challenges in SV analysis, particularly in handling ambiguous BND annotations, enabling flexible set operations, and automating complex analytical workflows. By unifying BND correction and flexible merging within a reproducible Snakemake pipeline, our tools streamline the transition from raw reads to high-confidence SV callsets, improving consistency across diverse platforms and samples. The modular design of our toolkit allows researchers to tailor analyses to specific biological questions, from population genetics to cancer genomics, while maintaining analytical rigor and reproducibility. We anticipate that these features will facilitate more reliable identification of SVs in population-level genomics, clinical diagnostics, and precision medicine research [38], ultimately advancing our understanding of the role of structural variation in human health and disease.

## Methods

### Software overview and implementation

OctopusV was implemented in Python and deployed through the Python Package Index (PyPI) for streamlined installation. The software provides a command-line interface with multiple subcommands, each addressing a specific aspect of SV (SV) analysis. Core functionalities include BND correction ("convert"), SV merging ("merge"), and benchmarking ("benchmark"), with all modules operating on customized VCF formats [15] to ensure compatibility with existing bioinformatics workflows. The software architecture comprises three interconnected layers: (i) the input processing layer for handling diverse VCF formats, (ii) the correction module for standardizing breakend representations, and (iii) the merging module for integrating multi-caller outputs through flexible set operations.

To further simplify real-world usage, we developed TentacleSV, a Snakemake-based automation layer built on top of OctopusV. TentacleSV coordinates the complete analysis pipeline from raw sequencing data through mapping, variant calling with user-selected SV callers, and finally OctopusV’s correction and merging operations. Users can customize their analysis through a minimal configuration file (config.yaml) that specifies input data locations, desired SV callers, and merging parameters. While OctopusV remains the core engine for BND correction and variant integration, TentacleSV ensures reproducibility by consolidating all analysis steps into one automated workflow, making it suitable for large cohort studies or multi-platform sequencing projects.

### BND correction algorithm

To standardize ambiguous breakend (BND) annotations, OctopusV employs a hierarchical classification system converting BNDs into three canonical SV types: INV, DUP, TRA. The algorithm initially classifies SVs into BND or non-BND events (**Fig. 3c**). BNDs are further divided into same-chromosome and different-chromosome types. Same-chromosome BNDs are analyzed through bracket orientation patterns in the ALT field to determine strand orientations, categorizing them as INVs (opposite orientations), DUPs (overlapping regions), or forward/reverse TRAs. Different-chromosome BNDs undergo mate-pair identification using positional coordinates specified in ALT fields, allowing a configurable positional tolerance (default: 3 bp). Mate pairs are classified into reciprocal TRAs (balanced reciprocal exchanges), merging TRAs (multiple breakpoints representing the same event), or independent TRAs (isolated events). Events lacking mate pairs are classified as special no-mate pairs—further analyzed based on positional relationships—or directly labeled as independent TRAs. All classified BND events are reformatted with standardized SVTYPE annotations, precise genomic coordinates, and orientation metadata, facilitating downstream interpretation. Detailed conversion rules are provided in **Supplementary Dataset 3**.

### SV merging and set operations

Following BND correction, OctopusV applies a systematic approach to merge SV calls from multiple callers. First, variants are sorted by their SV type and organized by chromosome pairs for efficient processing. The merging method differs between TRA and non-TRA events. For non-TRA variants (DEL, DUP, INV, INS), the algorithm groups events based on their genomic positions using adjustable distance thresholds (default 50–150 bp), while also considering size similarity (maximum ratio of 1.3) and overlap extent (minimum Jaccard index of 0.7). For TRA events, the algorithm uses more flexible position matching criteria and verifies that strand orientations are consistent between events before merging.

OctopusV offers various set operations for integrating results from multiple callers. Basic operations include union (keeping all unique variants), intersection (retaining only variants found by all callers), and threshold-based filtering where variants must be supported by a minimum number of callers (e.g., ≥2 or ≥3). Advanced operations unique to OctopusV include maximum support thresholds (e.g., variants supported by ≤2 callers), custom set operations that combine results using logical operators, e.g., ((Manta AND SVABA) NOT (Lumpy OR Delly)), and single-caller extractions for analyzing results from specific tools. These flexible set operations enable precise control over which variants are retained in the final output.

### Built-in benchmarking

OctopusV includes a benchmarking module for evaluating SV calls against truth sets or high-confidence callsets, following Genome in a Bottle (GIAB) consortium guidelines [11] with modified parameters. By default, it implements a matching algorithm that considers multiple criteria: position-based matching with a distance threshold of 500 bp, size similarity requirement of minimum 0.7 ratio, and variant type matching. These thresholds, which have been widely validated in large-scale benchmarking studies [42], can be adjusted by users to accommodate different analysis requirements or the specific characteristics of different variant types. The module calculates precision, recall, and F1 scores from true positives, false positives, and false negatives, both globally and stratified by SV type. Results are output as VCF files for each category (TP, FP, FN) along with summary statistics in JSON format.

### Data preparation

Simulated and real datasets were prepared to thoroughly assess OctopusV’s performance across SV types and sequencing platforms. For simulated data, VISOR (v1.1.2) was used to embed known SVs into the GRCh38 reference genome, sourced from dbVAR callsets, including CHM1 (nstd137) [43] and KWS1 (nstd106) [44], comprising 6,167 DELs, 9,899 INSs, 44 INVs, 3,712 DUPs, and 380 TRAs. Variants were integrated into two haplotypes using VISOR HAck with default parameters, mimicking a 30:70 ratio of homozygous to heterozygous events, followed by in silico sequencing at 30× coverage for short-read NGS, Oxford Nanopore (ONT), and PacBio platforms, incorporating platform-specific error profiles and coverage distributions. The ground truth for simulated datasets was constructed by merging variants from CHM1 and KWS1 dbVAR datasets, pre-processed through a rigorous pipeline: VCF files were sorted using BCFtools sort (v1.17) [14], normalized against GRCh38.p13 using BCFtools norm, and merged with BCFtools concat, removing duplicates (-d both option).

For real data evaluation, the NA12878 (HG001) genome was analyzed using NIST NA12878 HG001 HiSeq 30× data and PacBio Sequel II CCS data with a mean read length of approximately 11 kb. We created a ground truth reference by combining SVs from two sources: the 1000 Genomes Project [45] and a long-read assembly-based NA12878 SV dataset [21]. We processed this truth set using the same pre-processing pipeline described for the simulated data.

### Genome mapping and SV calling

Platform-specific mapping strategies were applied to both simulated and real datasets. Short-read data were aligned to GRCh38 using BWA-MEM2 (v2.2.1) [46] with parameters "-t 12 -M -Y" and appropriate read group information. Long-read data were processed using minimap2 (v2.28-r1209) [47] with parameters "-ax map-ont" for ONT and "-ax map-hifi" for PacBio, supplemented by "--MD -t 8 -Y" and read group tags. Following alignment, multiple SV callers were employed to generate variant calls. Short-read data were analyzed using Manta (v1.6.0), LUMPY (v0.2.13), SvABA (v1.1.0), and DELLY (v1.3.1) with default settings. Long-read datasets were processed using CuteSV (v2.1.1) with parameters "-l 50 -L 5000000 -r 1000 -q 20 -s 3", SVIM (v2.0.0), PBSV (v2.10.0), Sniffles2 (v2.5), SVDSS (v2.0.0), and DeBreak with parameters "--min_support 2 -t 8 --rescue_large_ins --rescue_dup --poa". For PBSV, a two-step process involving discovery with human tandem repeat annotations followed by variant calling with default parameters was implemented, generating multiple VCF files per dataset, including numerous BND records, for subsequent OctopusV analysis.

### BND prevalence analysis and correction evaluation

Two analyses were conducted to examine the importance of BND correction in SV studies. First, to quantify the prevalence of BND annotations, all SV calls from multiple callers were analyzed across five datasets (NA12878 NGS, NA12878 PacBio, VISOR NGS, VISOR ONT, VISOR PacBio). For each caller, the proportion of BND-labeled events relative to total SV calls was calculated to determine how frequently these ambiguous annotations occur in typical analyses. Second, to assess whether these BND events represent true biological variants rather than technical artifacts, the percentage of BND events matching high-confidence reference variants was determined through coordinate-based comparison against established truth sets.

To evaluate the BND correction functionality of OctopusV, we extracted BND-labeled entries from each caller’s output VCF files based on " [" or "]" notations in the ALT field. These entries were processed using OctopusV’s "correct" subcommand with default 3 bp position tolerance. We then matched the corrected events against ground truth datasets (VISOR ground truth for simulated data, NA12878 reference for real data) using position tolerances of 50 bp for non-TRA events and 5,000 bp for TRA events. Critically, our accuracy assessment focused exclusively on BND events that matched entries in the ground truth dataset. For these matched events, we calculated correction accuracy as the percentage correctly converted to their true SV type (INV, DUP, or TRA). This approach specifically evaluated OctopusV’s ability to standardize BND annotations rather than assessing SV caller performance.

The distribution of corrected BND events across final SV types (INV, DUP, TRA) was analyzed to characterize conversion patterns. For each SV caller on all three sequencing platforms (NGS, PacBio, ONT), the proportion of BND events converted to each SV type was calculated and expressed as percentages. The analysis included results from short-read callers and long-read callers.

To validate the biological significance of BND correction, selected events from NA12878 NGS data that were initially reported as BND by Manta and subsequently corrected by OctopusV were further examined. First, Truvari (v4.2.2) [42] was used to identify true positive SVs by comparing against the reference set. The validated variants were then annotated using AnnotSV (v3.4.4) with parameters "-SVminSize 0 -annotationMode full -overwrite 1" to identify events affecting functionally important genes. Selected events were then manually inspected using IGV to confirm the presence of supporting read evidence.

### Benchmarking of SV merging function

The merging performance of OctopusV was evaluated using five datasets: NA12878 NGS, NA12878 PacBio, VISOR NGS, VISOR ONT, and VISOR PacBio. Performance was compared against four widely-used SV merging tools: SURVIVOR (v1.07), Jasmine (v1.1.5), SVmerge (v0.36), and CombiSV (v2.3). Merging strategies evaluated included common operations (intersection, union, and minimum support thresholds) and OctopusV-specific operations (maximum support, caller-specific extraction, and custom set operations). VCF files generated by multiple SV callers were processed: short-read callers (Manta, Lumpy, SvABA, DELLY) for NGS datasets and long-read callers (CuteSV, PBSV, Sniffles, SVIM, SVDSS, DeBreak) for PacBio and ONT datasets. OctopusV merging was performed using the default parameters of the “merge” subcommand. SURVIVOR was configured with parameters “1000 1 1 1 0 30”, Jasmine with “--normalize_type” and four threads, and CombiSV and SVmerge used their default settings.

For NGS datasets (NA12878 NGS and VISOR NGS), merging strategies included intersection, union, minimum support (≥2 or ≥3 callers), maximum support (≤2 callers), caller-specific extraction, and custom set operations. Caller-specific extraction used Manta-specific variants due to Manta’s wide clinical use and its role as a standard component in many clinical pipelines. Custom set operations were defined as ((Manta AND SvABA) NOT (LUMPY OR DELLY)), as Manta [18] and SvABA [20] rely on local assembly, while LUMPY [19] and DELLY [27] primarily use read signals. For long-read datasets (NA12878 PacBio, VISOR PacBio, and VISOR ONT), additional minimum support levels (≥2 to ≥5 callers), maximum support, caller-specific extraction, and custom set operations were evaluated. Caller-specific extraction used Sniffles-specific variants, as Sniffles is a widely used long-read tool [48]. Custom set operations were set as ((DeBreak AND SVDSS) NOT Others), based on the shared partial order alignment (POA) method used by DeBreak [32] and SVDSS [31]. To enable comparison with CombiSV, which supports only a union operation with the caller set (Sniffles, PBSV, CuteSV, SVIM), an additional ‘4callers’ union analysis was performed across all NGS and long-read datasets.

OctopusV output files (SVCF: A customized VCF format for structural variants) were converted to standard VCF format using the “svcf2vcf” subcommand. SVCF is a customized version of the VCF format used by OctopusV for internal processing. This format keeps all standard VCF features while adding some fields needed for structural variants. It helps maintain consistent representation of corrected BND events and other SV types throughout the analysis. For more details about SVCF, see the OctopusV documentation (https://github.com/ylab-hi/OctopusV). Performance metrics (precision, recall, and F1 scores) were calculated using Truvari (v4.2.2) with default parameters (position matching thresholds: 50–500 bp, size similarity threshold ≥ 0.7, and --pctseq 0).

### Assessment of SV type consistency across merging tools

To evaluate the fidelity of SV type preservation during merging, we systematically assessed SV type consistency across five merging tools (OctopusV, SURVIVOR, Jasmine, SVmerge, and CombiSV) using datasets previously described (NA12878 NGS, NA12878 PacBio, VISOR NGS, VISOR ONT, and VISOR PacBio). We identified variants where the primary SVTYPE annotation was inconsistent with their constituent original variant types, generating a filtered VCF file containing incorrectly merged records. For each tool and dataset combination, the number of incorrectly merged variants was quantified. Detailed misclassification patterns, particularly prominent in SURVIVOR outputs, were further examined and visualized using Sankey diagrams. To evaluate the preservation of the original SV type distributions, proportions of each variant type (DEL, INS, DUP, INV, TRA) were calculated from true-positive calls generated by each merging tool and compared to the ground truth distributions. Results were summarized and visualized as normalized stacked bar plots.

## Declarations

### Ethics approval and consent to participate

Not applicable, as this study did not involve human participants, human data or human tissue.

### Consent for publication

Not applicable, as this manuscript does not contain data from any individual person.

### Availability of data and materials

The OctopusV and TentacleSV software packages are freely available at GitHub (https://github.com/ylab-hi/OctopusV; https://github.com/ylab-hi/TentacleSV) under the MIT license. All analysis scripts, configurations, and benchmarking results used in this paper are available at GitHub (https://github.com/qingxiangguo/OctopusV_paper). The GRCh38 reference genome was obtained from GENCODE (https://ftp.ebi.ac.uk/pub/databases/gencode/Gencode_human/release_38/GRCh38.p13.genome.fa.gz). For real data evaluation, we used the NA12878 NIST HG001 HiSeq subsampled 30X data (ftp://ftp-trace.ncbi.nlm.nih.gov/giab/ftp/data/NA12878/NIST_NA12878_HG001_HiSeq_300x) and PacBio SequelII CCS data from GIAB (https://ftp-trace.ncbi.nlm.nih.gov/giab/ftp/data/NA12878/PacBio_SequelII_CCS_11kb/HG001.SequelII.pbmm2.hs37d5.whatshap.haplotag.RTG.trio.bam). The ground truth sets used for read-data benchmarking include variants from the 1000 Genomes Project (ftp://ftp.1000genomes.ebi.ac.uk/vol1/ftp/phase3/integrated_sv_map/ALL.wgs.mergedSV.v8.20130502.svs.genotypes.vcf.gz) and the long-read assembly based NA12878 reference SV dataset (https://github.com/stat-lab/EvalSVcallers/blob/master/Ref_SV/NA12878_DGV-2016_LR-assembly.vcf) [21]. The simulated SV callsets were generated using variants from dbVAR, including CHM1 (ftp://ftp.ncbi.nlm.nih.gov/pub/dbVar/data/Homo_sapiens/by_study/vcf/nstd137.GRCh37.variant_call.vcf.gz) and KWS1 (https://ftp.ncbi.nlm.nih.gov/pub/dbVar/data/Homo_sapiens/by_study/vcf/nstd137.GRCh38.variant_call.vcf.gz) datasets.

### Authors’ contributions

QG and YL designed and implemented the core algorithms and computational framework. QG, YL and TYW conceptualized the project and designed the study. QG and YL performed the benchmarking analysis and validation experiments. TYW and AR contributed to project planning and provided critical feedback on tool development. QG, YL and RY wrote the manuscript with input from all authors. RY supervised the project.

## Acknowledgements

This project was supported in part by NIH grants R35GM142441 and R01CA259388 awarded to RY.

