## Supplementary Fig. 1, Supplementary Fig. 2, Supplementary Fig. 3 for "OctopusV and TentacleSV: a one-stop toolkit for multi-sample, cross-platform structural variant comparison and analysis"

**Supplementary Figure**

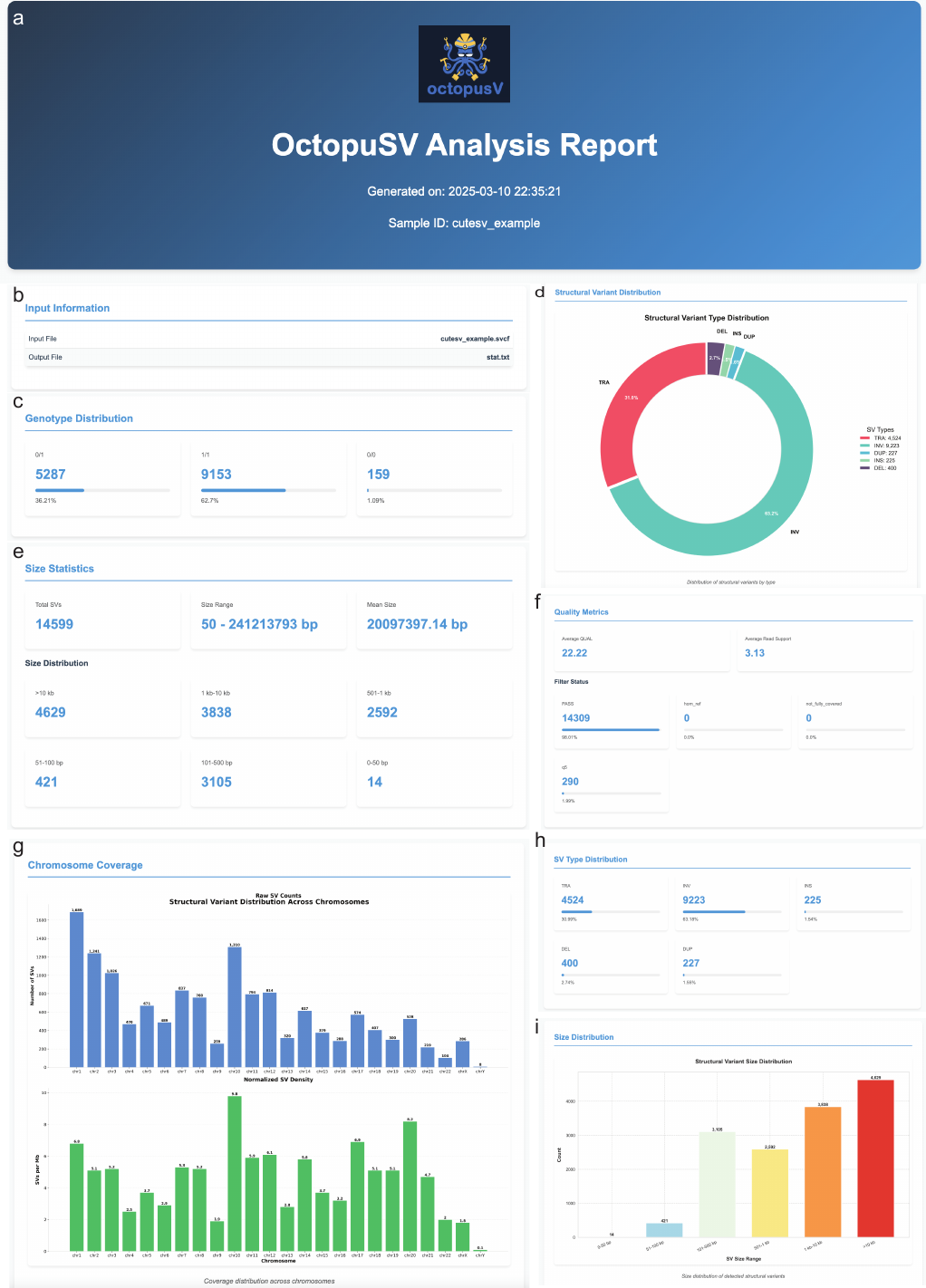

**Supplementary Figure 1.** **HTML report from OctopusV's statistical analysis module.** This figure displays the key visualizations generated by OctopusV's reporting system. Panels show essential structural variant metrics: **a-b** sample information, **c** genotype frequencies, **d, h** variant type composition, **e, i** size distributions, **f** quality metrics, and **g** chromosomal distribution. The HTML format allows researchers to interact with the data for efficient exploration of complex SV profiles.

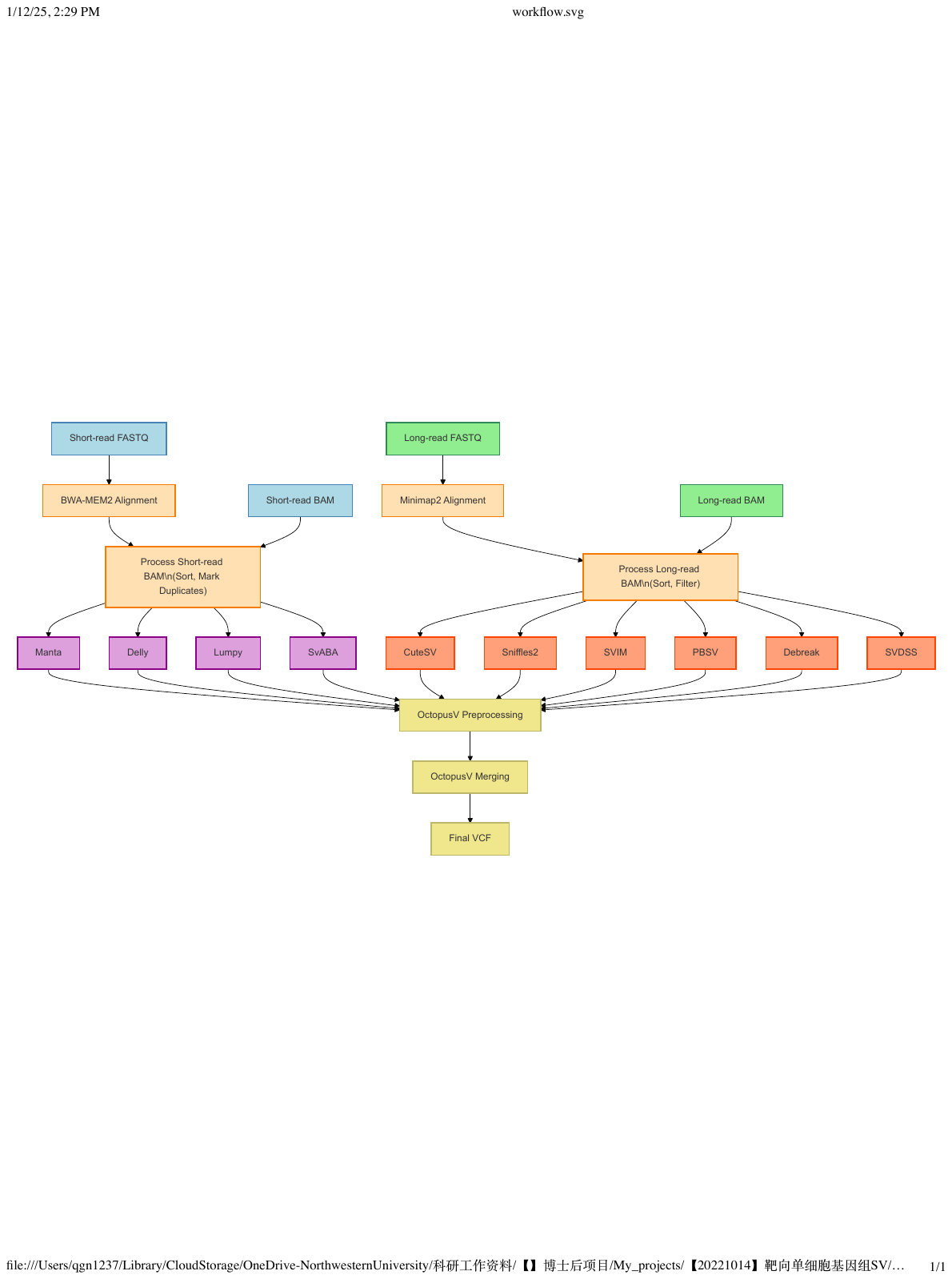

**Supplementary Figure 2.** **Workflow diagram of TentacleSV pipeline.** It processes both short-read and long-read sequencing data (FASTQ/BAM) through multiple SV callers and merges the results using OctopusV to generate a final VCF output.

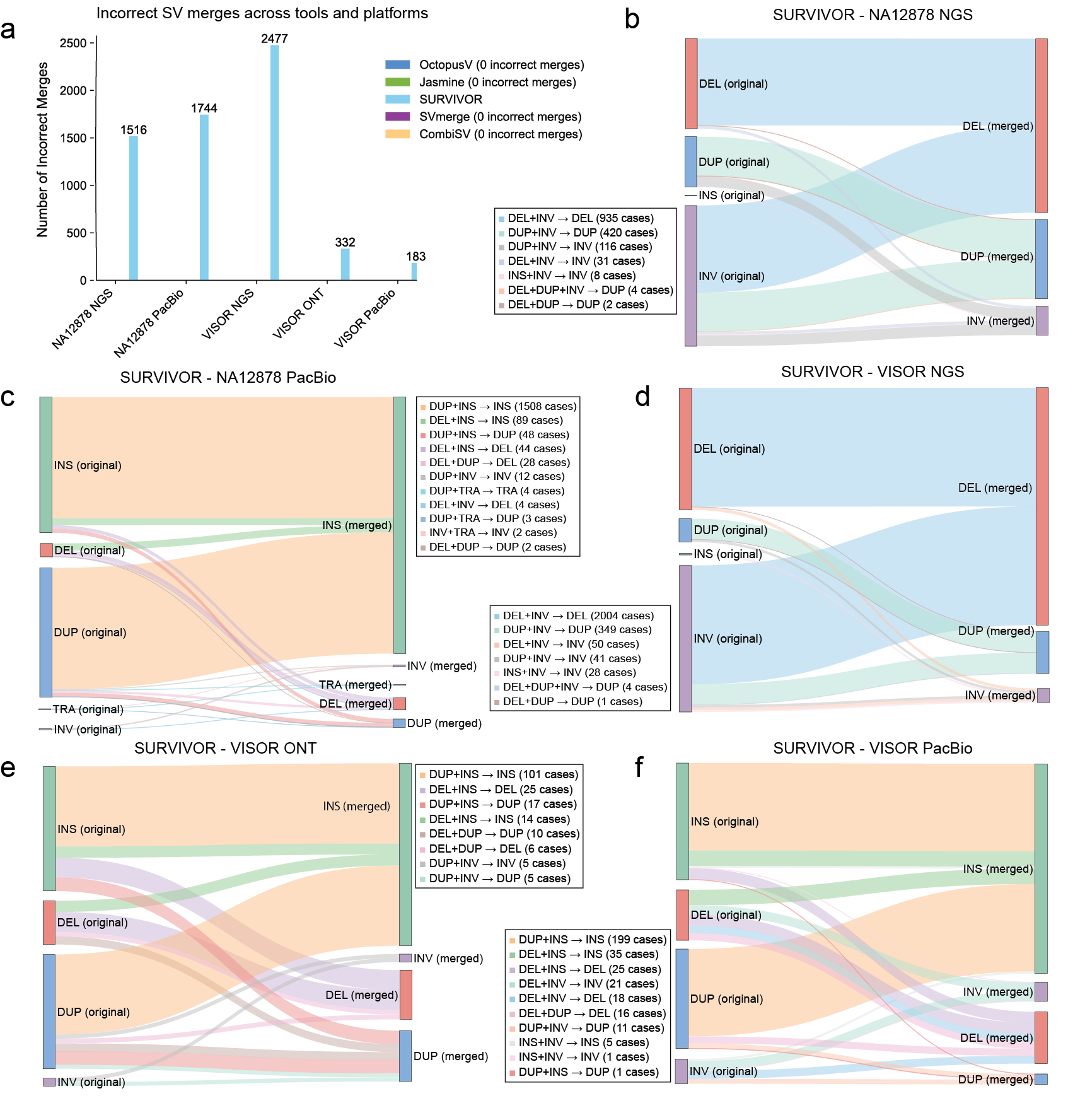

**Supplementary Figure 3.** **Evaluation of SV type consistency during merging across different tools.** **a** Comparison of SV type concordance across five datasets, showing varying degrees of type preservation among different tools. **b-f** Sankey diagrams visualizing SV type transitions during the merging process for SURVIVOR across different datasets: **b** NA12878 NGS, **c** NA12878 PacBio, **d** VISOR NGS, **e** VISOR ONT, and **f** VISOR PacBio. Flow widths represent frequency of specific type transitions, with annotation boxes providing counts of each transition pattern.
